# Basin-scale dynamics and enrichment-enabled genomics of marine nitrifiers: seasonality, niches, interactions, and genomic uniqueness

**DOI:** 10.1101/2025.02.05.636653

**Authors:** Sookyoung Kim, Elisa D’Agostino, David M. Needham

## Abstract

Nitrification occurs widely from the deep sea to animal holobionts, but the eco-evolutionary forces shaping the niches and dynamics of the lineages of the chemoautotrophic bacteria and archaea responsible remain largely unknown. To make strides towards this goal in a rapidly changing, exemplar marine ecosystem, the Baltic Sea, we studied basin-scale nitrifier spatio-temporal dynamics, coupled with enrichment-enabled comparative genomics. Based on metagenomes and rRNA gene sequencing, we found nitrifiers to be persistently relatively abundant throughout deep depths (>25 m), and from late-fall to spring in surface waters, as revealed by twice-weekly sampling across two years in the southwest Baltic Sea surface waters. In these surface waters, we observed time-lagged dynamics between ammonia- and nitrite-oxidizers, which were positively correlated with nitrite, nitrate, and diverse other prokaryotes, and negatively correlated with day length, light, and chlorophyll. For the dominant nitrifiers, ammonia-oxidizing archaea (AOA), we enriched five novel species including the dominant deep Baltic Sea species, and obtained genomes from all dominant AOA phylotypes. Among these genomes, which enabled fine-scale niche-differentiation, we observed a high degree of gene conservation, with most differences related to genes associated with interactions with the external environment, including genes involved in signal transduction, cell wall/membrane biogenesis, and inorganic ion transport, indicating these may be the primary drivers of strain-variability. We also observed differences in nitrogen and phosphorus metabolism between two dominant surface types. Together our study provides key insights into the niche of nitrifiers, and begins the process of understanding the mechanisms and functional implications of these patterns.

## INTRODUCTION

Nitrification is the microbial process whereby reduced nitrogen substrates are oxidized to nitrate. Nitrification is ubiquitous across natural (and artificial) ecosystems, and, in the deep ocean, the microbes involved are among the dominant carbon fixers [1–5]. Nitrification is a two-step process where ammonia is first oxidized to nitrite, and then, nitrate. In the marine environment, this process is carried out by a remarkably small number of families, with ammonia oxidation performed by archaea (AOA, [6, 7]) and bacteria (AOB, e.g., *Nitrosomonas* and *Nitrosococcus*), and nitrite oxidation by bacteria (NOB, e.g., *Nitrospina*, *Nitrospira*)[8, 9]. Some *Nitrospira* species are capable of comammox (COMplete AMMonia OXidation) - the oxidation of both ammonia and nitrite [10, 11]. While these comammox organisms have not yet been found to be abundant in the ocean, they have been observed in estuarine and brackish environments [12]. In addition to their role in nitrification, nitrifiers are often the numerically dominant carbon-fixing organisms in the deep sea, making them key players in the ecosystem [5, 13, 14]. Furthermore, ammonia-oxidizing archaea (Nitrosophaerota) have been proposed to be the model taxon for deep ocean prokaryotes due to their high abundance and ecological significance [15]. This highlights their importance in advancing our general understanding of microbial biology in the deep sea.

Within and between each nitrifier family, there is remarkable diversity and variation in the distribution and niches of taxa. While a combination of physiological, ecological, and evolutionary forces governs their distributions and dynamics, these factors are still poorly resolved for the marine environment. Contributing to the differences between families are some long-recognized traits [16–18]. For example, it is thought that AOA have higher affinity ammonia monooxygenase (AMO) enzymes compared to AOB, thus driving niche differentiation [19, 20] and allowing them to outcompete AOB in environments with low ammonia availability, and vice-versa. There is also trait variability within families - different AOA strains have diverse capabilities to take up alternative nitrogen substrates such as urea and cyanate [21–23], degrees of photoinhibition [18, 24, 25], and motility [26–28], among other factors [17]. Together these studies emphasize that the versatility of these populations may have important ecological implications in changing environments. Meanwhile, experimental evidence supports the idea that nitrifiers release carbon into the environment [29], which is ultimately taken up by co-occurring bacteria [29, 30] thus suggesting an important role in shaping ecosystem function.

The Baltic Sea presents an opportunity to study the ecology of nitrifiers in a natural ecosystem with a high environmental context that spans a spectrum of physiochemical conditions, which has led to it being considered a ‘time machine’ for ocean research [31]. In particular, this basin spans a full range of salinity, oxygen concentrations, and trophic states. It has been shown that nitrifiers are most active and abundant beneath the photic zone, in the redox-cline where oxygen is still available [32–34], but they are also present in sub-oxic, deeper depths [35–38], with a single AOA, identified as 16S rRNA clone sequence GD2, numerically dominating the redox-cline [34, 36]. While the presence of nitrifiers and nitrification has also been observed in the photic zone [39], time-series dynamics of nitrifiers in the surface waters have not been systematically analyzed.

Here, we utilized basin-scale (Baltic Sea) distributions and time-series sampling to advance knowledge of the niches of nitrifiers at fine phylogenetic resolution. We enriched five novel AOA species, including GD2, the dominant deep-sea strain, and obtained genomes from all dominant AOA, allowing us to explore their eco-genomics.

## MATERIALS AND METHODS

### Sample collection

Spatiotemporal data from the Baltic Sea was collected from across the Baltic Sea. Twice-weekly, depth-integrated seawater sampling was carried out for one and a half years from the Southwest Baltic Sea Kiel Fjord (KF, 54°19’48.3“N 10°09’01.9”E) and Boknis Eck time series (BE, 54°30’46.1” N 10°01’20.5” E). KF seawater (Oct 2021 to May 2023) was collected by bucket and processed within 2 hours. Temperature, oxygen, and salinity were measured continuously via the Kiel Marine Organism Culture Centre (KIMOCC; https://shorturl.at/bekTW) and temperature and salinity additionally via a handheld sensor (Conductivity portable meter ProfiLine Cond 3110 with Special conductivity measuring cells TetraCon® 325 S, WTW), which was used to support KIMOCC data, but not typically used. The full dataset is available via Supplementary Data 1. Weather data were taken from GEOMAR Meteorological Station (https://www.geomar.de/en/service/weather). At Boknis Eck, samples were collected monthly (Jan 2022 to June 2023) from 1 m, 10 m, 20 m, and 25 m. From the Broader Baltic Sea, we collected samples for amplicon sequencing from the Bornholm Basin (5 September 2022, 55° 37.50’ N, 15° 15’ 03” E) and Gotland basin (3 September 2022, 56° 54.90’ N, 19° 34.88’ E) via CTD/Rosette. Metagenomic samples collected in this study are listed in Supplementary Data 2.

For all locations, seawater was frozen for nutrient analysis at −20 °C until the run in the central analytical laboratory at GEOMAR using a standard colorimetric, Continuous Flow Analyzer [40]. 200 mL was filtered for chlorophyll on GF/F filters and extracted following with 90% acetone and read via fluorometry [40].

### Filtration, DNA extraction, and sequencing environmental samples

Seawater samples were vacuum-filtered under less than 200 mbar pressure through a 0.2 μm pore size filter (Pall, Supor™ 200 Membrane Disc Filters) and then stored at −80 °C until DNA extraction. DNA was extracted from each filter using the DNeasy Plant Kit (Qiagen) with a modified lysis protocol. Briefly, filters were thawed and transferred into screw cap microtubes with mixed glass beads (0.1 mm and 0.5 mm). Lysis buffer from the kit (AP1) was added, followed by three freeze-thaw cycles using liquid nitrogen and a 65 °C water bath. Mechanical lysis was conducted via bead-beating using a Laboratory Mixer (MM 400, Retsch). After lysis, lysate was transferred to a 96-well plate for purification via AcroPrep plate. DNA was eluted in Tris-EDTA and stored at −80 °C until amplicon or metagenomic library prep.

For amplicon libraries, 1 ng of DNA (quantified by Quant-iT PicoGreen dsDNA Assay Kit) was amplified and prepared for sequencing in single PCR reactions with universal V4-V5 primers 515F and 926R [41]. The forward primer included an Illumina P5 adaptor sequence (AATGATACGGCGACCACCGAGATCTACAC), a unique 8-base index with every index differing by at least two base pairs, a forward sequencing primer (ACACTCTTTCCCTACACGACGCTCTTCCGATCT), a random spacer of 7 random bases (NNNNNNN) to enhance diversity for the initial base-pair reads on the Illumina MiSeq. The reverse primer was similarly constructed but with P5 replaced with P7 (CAAGCAGAAGACGGCATACGAGAT) and reverse sequencing primer replaced with (GTGACTGGAGTTCAGACGTGTGCTCTTCCGATCT). PCR was performed with the KAPA HiFi HotStart PCR Kit (Cat. KK2102, Roche), using 0.4 U of KAPA HiFi HotStart DNA Polymerase, 1× KAPA Hifi Fidelity buffer, 0.3 mM dNTPs, and 0.4 μM of each primer. PCR thermocycling consisted of initial denaturation at 95 °C for 5 m, followed by 30 cycles of denaturation at 98 °C for 20 s, annealing at 55 °C for 15 s, and extension at 72 °C for 1 m, and a final extension at 72 °C for 5 m. Products were cleaned and size-selected via 0.8× (vol: vol) Ampure XP magnetic beads (Agencourt® AMPure® XP, Beckman Coulter), or normalized with Charm Biotech Just-a-Plate 96 PCR Normalization plates. For Ampure cleaned samples, DNA concentration was quantified with Quant-iT PicoGreen dsDNA Assay Kit and pooled in equimolar concentration. Prior to sequencing, the pooled DNA was purified again using 0.8× (vol: vol) of the magnetic beads.

For metagenomic libraries, 1 ng of DNA was prepared with Hackflex [42], a modification of the protocol of Illumina DNA Prep (M) Tagmentation Library. Following Gaio et al, DNA was tagmented in 50 µL reactions and then amplified with Phusion® Hot Start Flex DNA Polymerase (New England Biolabs, M0535) and amplified with a forward and reverse primers that consisting of P5 or P7 adapters, 8-base-indexes, and partial overhangs to the transposome adapter TCGTCGGCAGCGTC or GTCTCGTGGGCTCGG, respectively. PCR was carried out with an initial denaturation at 98 °C for 30 seconds, followed by 12-16 cycles of: 98 °C for 10 s, 62 °C for 30 s, 72 °C for 30 s, and 72 °C for 5 m for a final extension. After PCR, products underwent 0.6× size selection via Ampure XP magnetic beads. The products were then re-evaluated using a bioanalyzer, followed by an additional 0.8x bead clean-up to remove short DNA fragments.

All sequencing was performed at the Competence Centre for Genomic Analysis (CCGA) Kiel, with amplicon libraries sequenced on MiSeq (2×300 bp) and metagenomic libraries on Novaseq (2×150 bp).

### Enrichment and sequencing of AOA

Archaeal medium was adapted from [43], with modifications. In brief, surface water was collected from the Baltic Sea (Kiel Fjord) and then aged. This water serves as the base of the medium. Cultures were initially established using 0.2 µm filtered but non-autoclaved seawater (with the exception of *Ca.* N. eckernfoerdensis and *Ca.* N. balticoprofundus) (to avoid potential adverse effects on organic content) and amended with 100 μmol l−1 NH4Cl, 1 ml l−1 chelated trace element solution, 15 μmol l−1 KH2PO4 and 50 μg ml−1 ampicillin. After obtaining stable AOA enrichments with consistent nitrite production, subsequent transfers were maintained in autoclaved seawater media with the same amendments. To reduce the diversity and abundance of bacteria, two selected strains were further supplemented with 50 μg/ml of streptomycin and kanamycin. The enrichment cultures were maintained at room temperature, except for the dominant deep Baltic Sea strain (N. balticoprofundus) which was enriched at 10°C.

Growth was monitored through nitrite accumulation, indicating AOA activity, and 10% volume of enrichment was transferred once reaching 70 μM NO2. For DNA extraction, 1-2 mL culture was filtered through a 0.2 μm Supor filter at <200 mbar pressure, stored at −80 °C, followed by the same modified lysis protocol we used for the environmental samples, and then following manufacturer’s instructions for purification via the DNeasy Plant Kit (Qiagen). For rRNA gene amplicon sequencing library preparation, we followed the same protocol described for environmental samples but used 1 µL of DNA extract with 30 PCR cycles. Enrichment cultures showing distinct ASVs are selected for metagenomic sequencing using the Hackflex protocol (12-16 PCR cycles).

### Amplicon and metagenomic data processing

For rRNA gene amplicon data, the primers 515F and 926R were trimmed from raw reads using Cutadapt v4.4 [44]. 16S and 18S amplicons were split with bbsplit.sh [45] by comparison to SILVA 132 [46] and PR2 [47]. The subsequent analysis focused exclusively on prokaryotic reads, using QIIME2 v2022-2 and/or v2022-11 [48]. Forward and reverse reads were based on sequence quality scores from FastQC v0.12.1 [49] and denoised to amplicon sequence variants (ASVs) with DADA2 [50]. Taxonomic classification was performed on ASVs with vsearch [51] and the SILVA 138.1 database [46]. Individual sequencing runs were analyzed separately and subsequently merged as recommended, [50]. Chloroplast, mitochondrial, and unassigned reads were then removed using the taxa filter-table function in QIIME2. Reads originating from the spiked-in controls were manually identified and removed. Relative abundance of each sample was calculated by division by the total number of sequences remaining prokaryotic sequences in each sample. Samples with low sequencing depth (<900 reads) were filtered out.

For metagenomic analysis, paired-end reads were quality filtered with Trimmomatic v0.39 [52] by truncating at the first base with a quality score below three, followed by further truncation if the moving average quality score across 25 base pairs fell below 30 (settings: LEADING:3 TRAILING:3 SLIDINGWINDOW:25:30 MINLEN:50). The resulting paired reads were then assembled with SPAdes v3.13.0 [53], with k-mer sizes (21, 33, 55, 77, 99, 127) and with ‘-sc’ option. For downstream analysis, only contigs exceeding 5,000 base pairs were retained. Quality filtered paired reads were mapped back to contigs Bowtie2 v2.3.5 [54] for each respective sample. Enrichments were processed individually. All sets of metagenomic reads were mapped to the individual sample assemblies, to enable binning while reducing the potential of chimeric contigs. For enrichments, the binning process was conducted using the Anvi’o v8 interactive interface [55–57], leveraging information such as tetranucleotide frequency, GC-content, and contig coverage. For environmental samples, the data was first automatically binned with Metabat and then refined using Anvi’o. Environmental metagenomic bins were then dereplicated with dRep. The quality of all resulting bins from enrichment and environmental samples was evaluated using CheckM v1.2.2 (Parks et al., 2015). Finally, taxonomic classification for all bins was performed using GTDB-TK v2.1.1 (Chaumeil et al., 2019).

### 16S rRNA gene phylogenetics and phylotyping

rRNA gene reads were extracted via SortMeRNA (-e 0.0001) from 15 metagenomes first reported here as well as from 169 public metagenomes [58–61](Supplementary Data 2), based on the smr_v4.3_sensitive_db.fasta database. The taxonomic classification of the reads was used to select *Nitrosopumilus*, Nitrospinaceae, Nitrospiraceae, Nitrosomonadaceae, and Nitrosococcaceae. Relative abundances were calculated from the number of reads classified as these respective groups divided by the total amount of reads predicted to be 16S rRNA gene reads in general.

In order to phylogenetically classify (phylotype) the 16S rRNA gene amplicons and 16S rRNA reads from metagenomes for the groups above, we constructed phylogenetic trees of the 16S for *Nitrosopumilus*, Nitrospinaceae, Nitrospiraceae, Nitrosomonadaceae, and Nitrosococcaceae. The phylogenetic trees were assembled from a variety of resources. First, we selected 16S rRNA genes from greengenes2 [62] database. Second, we assembled 16S rRNA gene reads from quality-filtered metagenomes derived from Baltic Sea samples with matam [63]. In the case of *Nitrosopumilus* we also extracted 16S rRNA reads from reference genomes as well as those assembled from our own data. In all cases, the 16S rRNA genes longer than 800 bp were clustered with vsearch at 0.995 sequence similarity, selecting the centroid as the representative sequences. Sequences were then aligned with muscle default settings, trimmed with trimAI [64](-gt 0.1), and phylogenetic trees constructed with IQ-TREE (-m GT −B 1000 –alrt 1000). The resulting trees were clustered by TreeCluster.py (-t 0.01, -s 0.80 -m max_clade) in order to group clades. ASV sequences as well as those predicted by SortMeRNA were then placed on these trees with epa-ng (--filter-min-lwr 0.01). Resulting jplace files were processed with gappa [65] and converted to csv with guppy.

### Phylogenomics and comparative genomics of AOA

*Nitrosopumilus* genomes considered as species representatives were retrieved from GTDB data v214.1 [66] using GToTree v1.8.1 [67], with *Nitrosopelagicus brevis* as an outgroup. Each reference genome underwent a quality assessment using CheckM v1.2.2 [68] to ensure a minimum of 80% completion and less than 5% redundancy. From the initial set of reference genomes, those showing >95% average nucleotide identity with genomes from this study were excluded (n=2), because in both cases they were of poorer quality in terms of genome completion and/or number of contigs. The final dataset encompassed 60 genomes, including 52 *Nitrosopumilus* representatives from GTDB, N. brevis as an outgroup and seven from this study (Supplementary Data 3). For incomplete genomes, estimated genome sizes were calculated using a combination of completion and redundancy estimates [69].

The phylogenomic tree was constructed by GToTree v1.8.1 [67], with the archaeal single-copy genes set (76 target genes). The analysis pipeline included gene prediction using Prodigal v2.6.3 [70], target gene identification with HMMER3 v3.3.2 [71], individual gene alignment using Muscle 5.1.linux64 [72] and alignment refinement with trimAl v1.4.rev15 [64] before concatenation. The phylogenetic tree was estimated using IQ-TREE [73–75], supported by 1000 bootstrap replicates. The tree was visualized with R 4.2.2 using ggtree v3.6.2 [76] and other tools to process tree data [77, 78]. For functional gene annotation, selected gene sets (amoABC, uvrABC, ectABC and thpD, pstABC, ureABCDEFG, cheCBRWY/flaB and 3-hydroxypropionate/4-hydroxybutyrate (3HP/4HB) cycle related genes [79] were manually identified from various *Nitrosopumilus* species, with gene IDs manually gathered from NCBI and UniProt (Supplementary Data 4). These genes were downloaded using the ‘ncbi datasets’ tool and analyzed using NCBI BLAST+ [80]. BLAST results were filtered (e-value >1e-5 and bitscore >100) and summarized in a gene presence-absence table. The presence of each functional category/pathway (Fig. 6) was determined based on whether more than half of the related genes were detected. Gene patterns and metadata are visualized using ggplot2 [81]. The alignment of these elements next to the tree was achieved using the aplot package.

To compare all the genomes within the genus *Nitrosopumilus*, we employed Anvi’o v8 [56, 57]. For each genome, databases were created and annotated with important gene sets (rRNA and single-copy core genes of taxa) using built-in databases, with initial functional assignments using the Clusters of Orthologous Genes (COGs) database [82]. Protein sequences were aligned using BLASTP [80] and clustered using MCL [83] with an inflation value of 10 for fine-scale cluster division. To analyze gene distribution patterns, we classified the gene clusters into four categories based on the number of genomes contributing genes to each cluster. Given the inclusion of metagenome-assembled genomes (MAGs) to pangenomics, we adopted more flexible classification criteria for gene clusters [84]: core (genes present in 51-60 genomes, ≥85%), semi-core (28-50 genomes, 47-83%), flexible (3-27 genomes, 5-45%), and rare (1-2 genomes, <5%). The gene cluster summary was exported and additional functional annotations were obtained using eggNOG-mapper [85]. These annotations were integrated with the cluster summary data to examine how genes from each COG20 functional category were distributed among core, semi-core, flexible, and rare gene cluster categories.

We used the 60 representative genomes as references to map the metagenomic data from at 0.95 similarity using bbmap. The ‘trimmed means’ were then computed with coverm genome default settings: --contig-end-exclusion and –min-covered-fraction of 0.25. Statistical analyses of environmental data

To analyze the correlation between the relative abundance of each microbe and environmental parameters, eLSA v1.0.2 [86] was used to calculate the time-lagged correlations using data from Kiel Fjord. Samples lacking 16S rRNA amplicon data or with low reads (<800) were excluded, and ASVs observed in more than 25% of samples were retained (i.e., at least 41 out of 161 samples). For environmental data, missing values were manually linearly interpolated within the time series and eLSA’s internal interpolation functions were not used. The analysis was run with a maximum time-lag of six sample points (∼three weeks), and significance values were calculated with the ‘mixed’ method using 1000 permutations. To analyze the correlation between abundant nitrifiers and environmental parameters, key ASVs and relevant environmental parameters were extracted from the eLSA results and visualized as a heatmap using R (v4.2.2) with ggplot2, with hierarchical clustering performed using Ward’s method. The resulting correlation network was filtered to retain only strong and significant correlations (correlation >0.7, p ≤0.001, q ≤0.001). The filtered network was visualized using Cytoscape, summarizing important nodes and edges associated with nitrifiers.

## RESULTS AND DISCUSSION

### Basin-scale distributions of nitrifiers

To gain an overview of the distributions and molecular diversity of nitrifiers in the Baltic Sea, we surveyed 15 new metagenomes and 169 published water-column metagenomes [58–61]. Based on 16S rRNA gene reads extracted from these metagenomes, this meta-analysis revealed that nitrifiers, in particular Nitrosopumilaceae, Nitrosomonadaceae, and Nitrospinaceae are common members of the microbial community across the entire Baltic Sea throughout the water column (Fig. 1). In the deeper depths, these three groups were almost always detected, but they were less commonly detected in surface waters (<25 m) (Fig. 1). While there was high variability between samples and depths, Nitrosopumilaceae was overall the most relatively abundant group of nitrifiers, making up 0.53% of prokaryotic rRNA gene reads (Fig. 1), followed by 0.12% for both Nitrosomonadaceae and Nitrospinaceae. Meanwhile, Nitrosococcaceae and Nitrospiraceae made up a more minor percentage of 0.009% and 0.003%, respectively (Supplementary Data 2). The lower abundance of these latter two groups led us to focus our analyses on Nitrosopumilaceae, Nitrosomonadaceae, and Nitrospinaceae due to their more robust recovery.

**Fig. 1:**
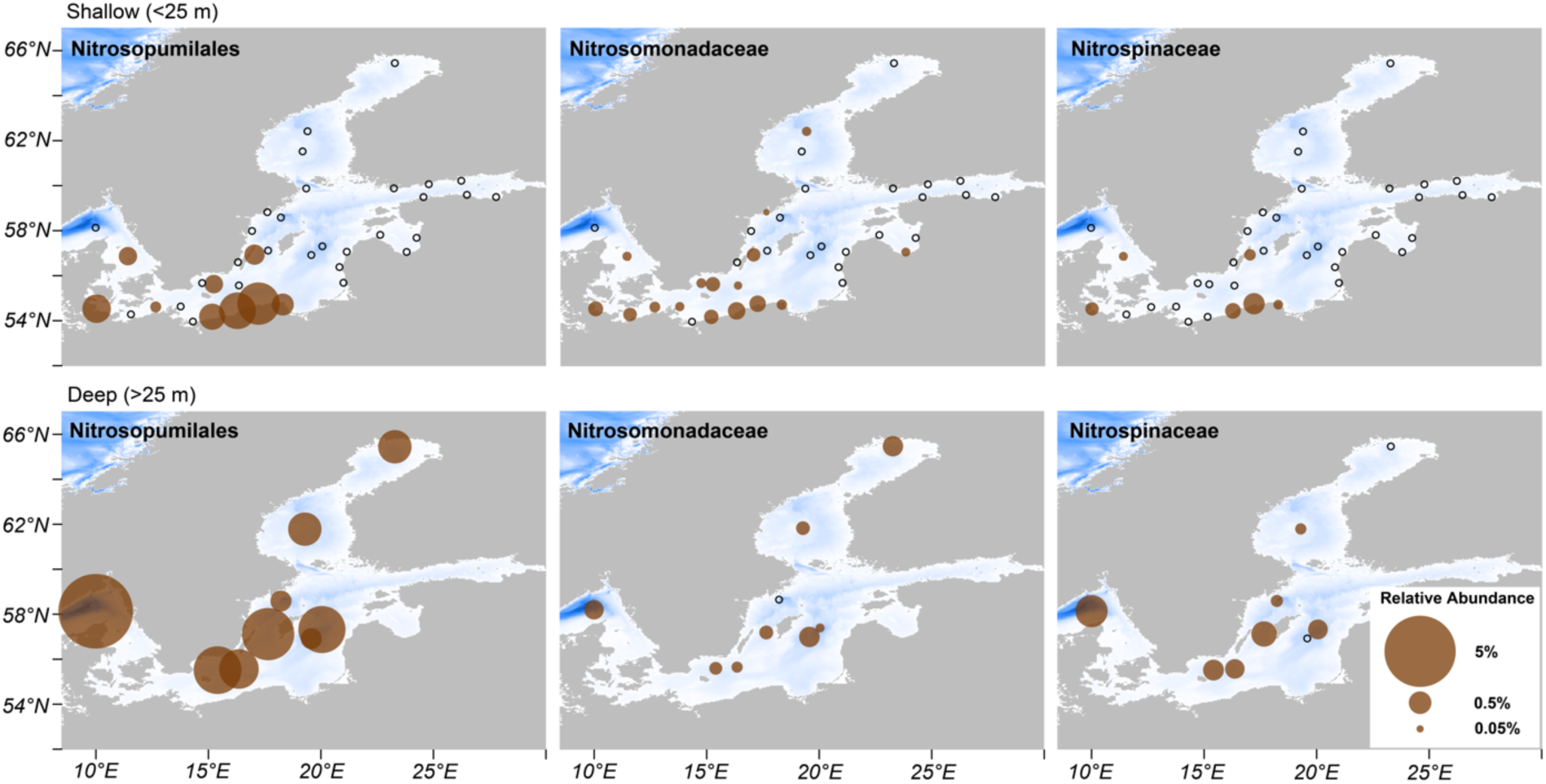
Distributions of nitrifiers across the entire Baltic Sea. Relative abundance of nitrifier rRNA gene sequences were extracted and classified with SortMeRNA. The pie size indicates the average relative abundance of the given family at each location (all samples within 1° of latitude and longitude are averaged). The top row are surface samples (0-25 m), and the second row are deep samples (>25 m). Open black circles indicate no detection. The background colors of the Baltic Sea (whites and blues) indicate water column depth. We chose this metagenomic approach as many rRNA amplicon datasets exclude Archaea, and reference genomes are limited for AOB and NOB. Note, AOA whole genome mapping results are presented in Supplementary Fig. 5.

Beyond the overall abundance patterns, we found that the predominant phylotypes of the nitrifiers differed in their abundances between surface and deeper depths, especially for AOA and NOB, but less so for AOB (Supplementary Fig. 1). For AOA, a single phylotype dominated the surface waters. This phylotype was related, among cultured AOA, to *N. limneticus* [87]. A second phylotype related to *N. oxyclinae* HCE1 [88] and GD2 [34] was more enriched in the deeper depths (Supplementary Fig. 1). For AOB, in surface waters, phylotypes related to the uncultivated genera GCA-2721545 and VFJL01 showed similar abundances, while GCA-2721545 was most relatively abundant in deep waters (Supplementary Fig. 1). For NOB, within the Nitrospinaceae family, we found depth-specific dominance of two genera: *Nitromaritima* dominated in surface waters, and LS-NOB in deep waters (Supplementary Fig. 1). In the case of both NOB and AOB, there were no isolates within the phylotypes of these sequences; the best hits were to rRNA genes extracted from Metagenome Assembled Genomes (MAGs).

To complement the basin-scale observations from metagenomes, we utilized universal rRNA gene amplicon primers from two depth profiles in the Baltic Sea (Supplementary Fig. 2). In deeper waters (>25 m), AOA were again dominant, with the most abundant AOA ASV coming from the same phylogenetic group that dominated the deep metagenomes, and this ASV is 99.7% similar to the rRNA gene of the GD2 (376 out of 377 bp across the ASV region) (Supplementary Fig. 3). We refer to this ASV as ASV N. balticoprofundus, as we successfully enriched and genome sequenced an AOA with the exact 16S rRNA gene sequence (*Ca.* N. balticoprofundus, reported in detail below). The other AOA ASVs that were less abundant are primarily affiliated with the AOA group found in the surface metagenomes. We also connected these ASVs to MAGs and refer to them as ASV BACL13 MAG-121220-bin23, and ASV N. kielensis (based on matches to bins of the same species, *Ca.* N. kielensis, reported in detail, below). From the amplicon dataset, AOB and NOB were also more relatively abundant in deeper depths, with AOB more abundant in the mid-waters (five ASVs detected), while NOB was more abundant in the deeper waters (11 ASVs detected, Supplementary Fig. 3).

Together, this basin-scale analysis revealed depth partitioning both in the overall abundances of nitrifiers, but also at the phylotype and ASV level. Given the only sporadic detection of nitrifiers in the surface waters, we set out to understand more about this variability.

### Seasonality of nitrifiers and environmental parameters

To understand the dynamics of nitrifiers in surface waters, we collected twice-weekly samples from the top meter of seawater in the southwest Baltic Sea between October 2021 and April 2023 (Kiel Fjord Time-series, Supplementary Fig. 2). Utilizing universal rRNA gene amplicon sequencing, we observed pronounced seasonality in nitrifier communities (Fig. 2). Among nitrifiers we found that AOA dominated in relative abundance, with ASV N. kielensis showing distinct seasonal patterns (Fig. 2A). This ASV peaked during October-November and maintained a high presence until March, with average relative abundance of 1.35% during October-March compared to 0.01% during April-September. ASV N. kielensis was 58-fold more abundant than the second most abundant ASV (herein, ASV N. decembris). Regarding interannual variation, overall AOA abundance was 3.7-fold higher in winter 2021-2022 compared to the following winter.

**Fig. 2:**
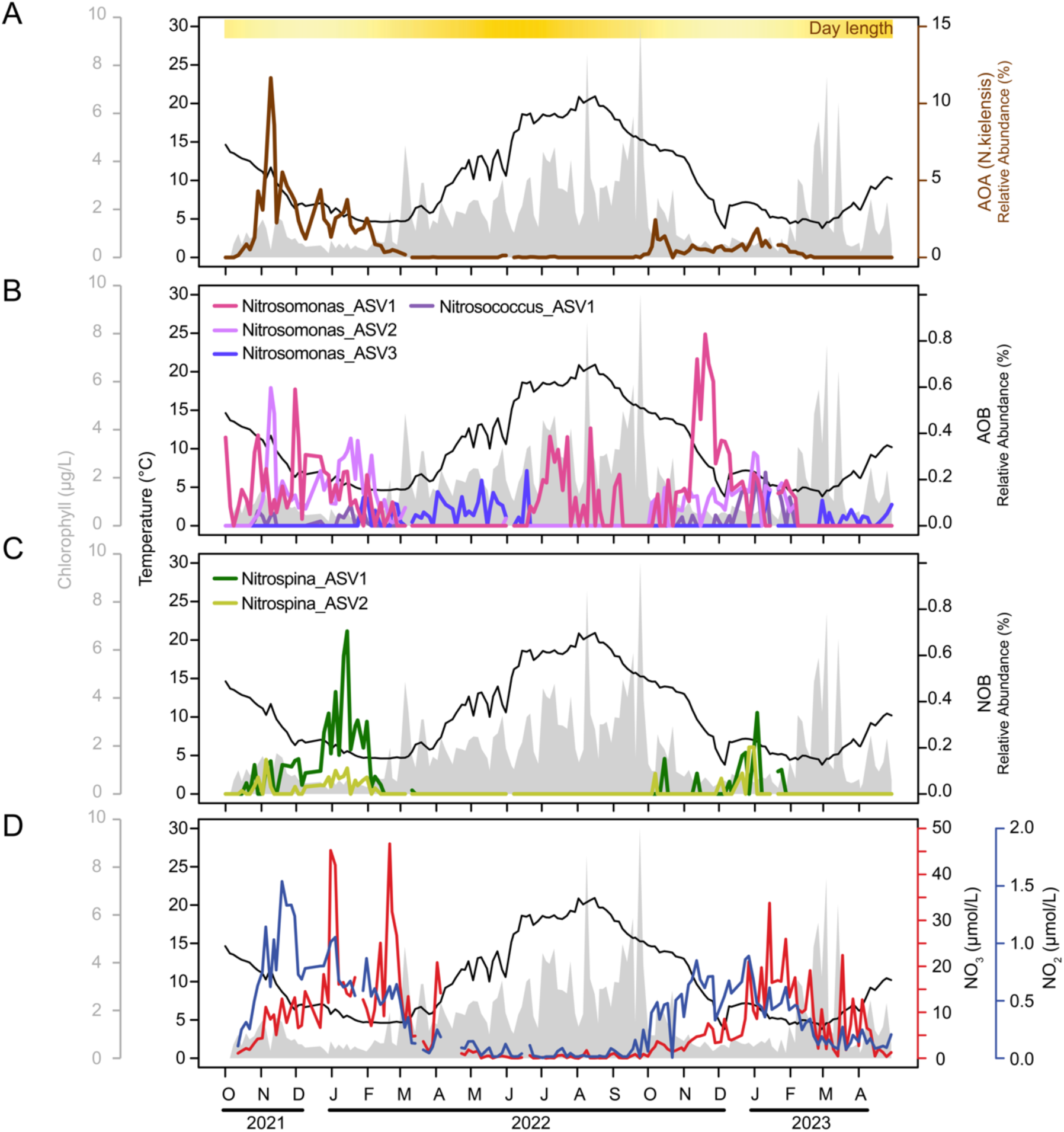
Time-series dynamics of nitrifier ASVs and inorganic nitrogen concentrations at the Kiel Fjord Time-series station based on 16S rRNA gene amplicon analysis. The top three panels show the relative abundances of A) AOA, B) AOB, and C) NOB ASVs, represented by colored lines. The yellow bar at the top of the first panel represents day length, with more intense yellow indicating longer days. The bottom panel displays NO₂ (blue) and NO₃ (red). Black lines indicate temperature, and gray shading shows chlorophyll concentration. A single AOA ASV (ASV N. kielensis) dominated the AOA community with 58-fold higher abundance than the second most dominant AOA ASV. AOA and AOB peaks occurred in late fall to early winter, followed by NOB peaks in mid-winter. The temporal dynamics of inorganic nitrogen species align with this succession, where NO₂ accumulates with ammonia oxidizer maxima and NO₃ with NOB peaks.

Beyond AOA, the other nitrifiers in the southwest Baltic Sea surface community were also more relatively abundant in the late-fall and winter months. However, unlike AOA in the surface, AOB were more diverse and periodically observed also in the summer, with a clear succession of three different AOB ASVs over time, each becoming the most relatively abundant ASV at different time points (Fig. 2B). NOB reached maximal relative abundance later in mid-winter, although it initially increased in abundance together with AOA and AOB. Similar to AOA, NOB were almost absent from spring to early fall (Fig. 2C).

We observed similar intra- and inter-annual patterns in nitrifier dynamics throughout the water column (1-25m) at Boknis Eck Time-series station (Supplementary Fig. 2), which we monitored monthly during the same period as the high-resolution time-series in Kiel Fjord. All the nitrifier groups had the highest relative abundances in late-fall and winter months, particularly in the deeper water (10-25 m), though they were also all detectable at 1 m, especially AOA and AOB (Supplementary Fig. 4). Unlike the surface waters, the deeper layers (20-25m) at Boknis Eck maintained a substantial nitrifier population even during some summer months.

Winter-time peaks in nitrifiers’ abundance, most notably among AOA, have been observed across various marine surface waters, including the northern Atlantic Ocean [89], Mediterranean Sea [90, 91], and in higher latitudes such as the Arctic and Antarctic oceans [92–94], and Barents Sea [93]. On the other hand, in subtropical coastal ecosystems such as southeastern USA (Sapelo Island) and southeast China (Zhejiang Province), their high abundances in summer were documented [95–97]. Notably, the seasonality we observed here has not yet been clearly documented in the Baltic Sea, likely because earlier regional studies used bacterial 16S rRNA primers that excluded AOA [98, 99]. The observation that AOA can have high relative abundances even in the surface is remarkable, given that ammonia oxidation is classically and experimentally shown to be inhibited by light [18, 24]. However, it seems apparent that they can overcome this limitation, when other factors are suitable for their success, although the mechanisms remain unknown.

### Correlations to environmental parameters

To understand the environmental factors potentially driving and being impacted by the seasonal surface dynamics more fully across the high-resolution time-series, we examined a variety of physico-chemical factors. The nitrifiers, in general, had strong negative correlations to temperature, chlorophyll, light, day length, oxygen, among others (Fig. 3). On the other hand, nitrifiers had strong positive correlations with inorganic nutrient concentrations (Fig. 3). The inorganic nitrogen species showed a sequential progression of transformation that corresponded to nitrifier dynamics: NO₂ peaked earlier in the late fall, coincident with highest abundances of AOA and AOB, while NO₃ increased subsequently during the peak of NOB, reflecting the conversion of NO₂ to NO₃ (Fig. 2D).

**Fig. 3:**
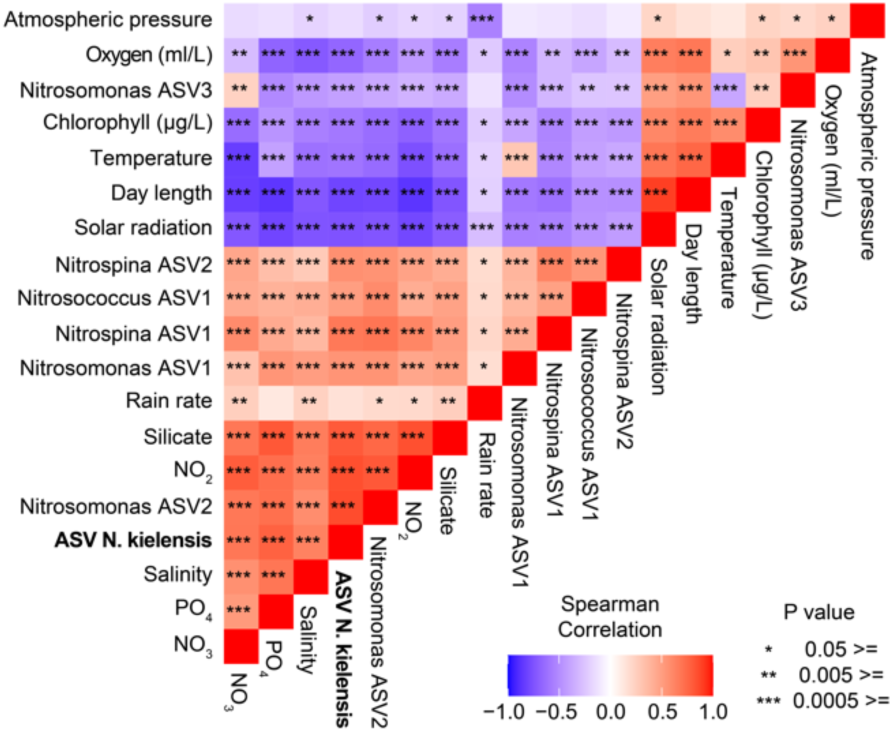
Heatmap showing the correlations between ASVs of nitrifiers (AOA, AOB, and NOB) and environmental parameters from the Kiel Fjord Time-series station over one- and-a-half years. Colors indicate correlation coefficients with asterisks denoting significance levels of correlations. The dominant southwest Baltic Sea surface AOA ASV (ASV N. kielensis) is highlighted in bold.

The exceptions to these general trends were two AOB ASVs that had peaks in warmer seasons. Nitrosomonas ASV1 had a positive correlation to temperature (peaking in summer), while Nitrosomonas ASV3 was positively correlated with solar radiation and day length (peaking in late spring-early summer) (Fig. 3). Despite their seasonal differences from other nitrifiers, both AOB maintained a positive correlation with nitrate. This suggests a niche differentiation where certain AOB demonstrate adaptability to high temperatures and compatibility with other groups occurring in the summer, contrasting with the winter-time preferences of AOA and NOB.

To further quantify the relationships of nitrifiers to environmental conditions in the twice-weekly temporal data, we conducted a time-lagged correlation network analysis to examine connections between nitrifiers, environmental parameters, and the broader microbial community (Fig. 4). Beyond the strong relationships between ammonia oxidizers and nitrite, we identified significant positive relationships with diverse prokaryotes from eight different orders, including prevalent community members such as SUP05, Cryomophaceae, NS4, SAR406, Planctomycetes, SAR324, and SAR11 Clade II. Notably, SAR324 was correlated with all three families of nitrifiers, showing a successional pattern with a time lag of ∼1.5 weeks after ammonia oxidizers and ∼2 weeks after NOB, suggesting either a delayed response to nitrification or a slower response to shared environmental triggers. Many of the taxa correlating with nitrifiers, particularly SUP05, SAR324, and SAR406, are typical deep-sea inhabitants with some representatives capable of carbon fixation, similar to nitrifiers. The sole negative correlation observed was between ASV N. kielensis and the summer-abundant Flavobacterium NS3a.

**Fig. 4:**
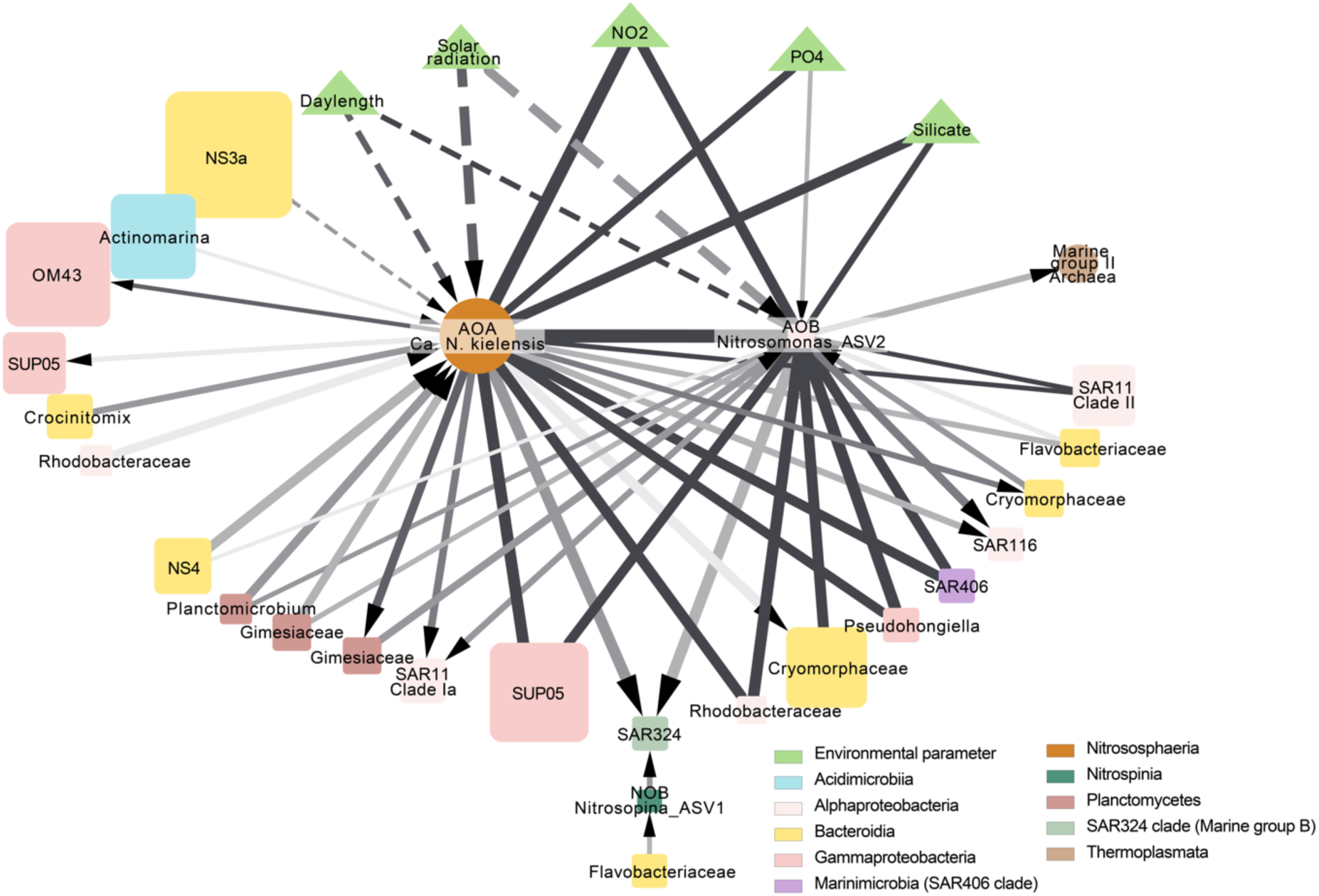
Network analysis of nitrifiers, associated microbial taxa, and environmental parameters from the Kiel Fjord Time-series station. Nodes represent taxa (squares: bacteria; circles: archaea) or environmental parameters (triangles), with node size proportional to mean relative abundance. Edges show significant time-shifted Spearman correlations (|ρ|>0.7, p ≤0.001, q ≤0.001) calculated with a lag tolerance up to three weeks. Edge color intensity reflects temporal relationships, with darker colors representing concurrent correlations and lighter colors indicating increasing time lags. Solid and dashed edges indicate positive and negative correlations, respectively. Arrows denote the direction of delayed responses.

### Novel enrichment and genome characterization of AOA

Our observations indicated AOA as the dominant nitrifier across temporal and spatial scales in the Baltic Sea, underscoring their ecological significance. Despite their importance, AOA genomes are generally challenging to recover from metagenomic and single-cell sequencing studies [100], and there was only one cultured representative from the Baltic Sea prior to our study [37]. Thus, to advance our understanding of these abundant nitrifiers, we set out to obtain representative genomes for the dominant AOA.

Through enrichment cultivation and subsequent metagenomic sequencing, we were able to generate five high-quality, near-complete genomes of novel *Nitrosopumilus* candidate species from the Baltic Sea, complementing 54 high-quality representatives in the GTDB database (v214.1). In these enrichments used for genome sequencing, AOA made up between 5% and 80% of the prokaryotes, with a diverse set of bacteria also inhabiting the enrichments (Fig. 5A). The genomes were obtained via manual curation of the assemblies based on tetra-nucleotide frequency, GC, and coverage, in which AOA genomes stand out starkly to the bacteria in the enrichments (Fig. 5B). Among the enriched AOA was the dominant deep Baltic Sea strain, which corresponds to GD2 (99.7% 16S rRNA similarity, 1315/1319 bp) and our ASV N. balticoprofundus (100% similarity). We enriched this candidate species from 63 m depth in the Bornholm Basin and designated it as *Ca.* Nitrosopumilus balticoprofundus. We also assigned *Candidatus* status to four additional novel candidate species, which were typically found in surface waters although relatively rare (See Protologues for new *Candidatus* species). This effort significantly improved the representation of Baltic Sea AOA, making it the marine environment with the most AOA enrichments (or cultures), followed by the Pacific Northwest, USA (Fig. 5C).

**Fig. 5:**
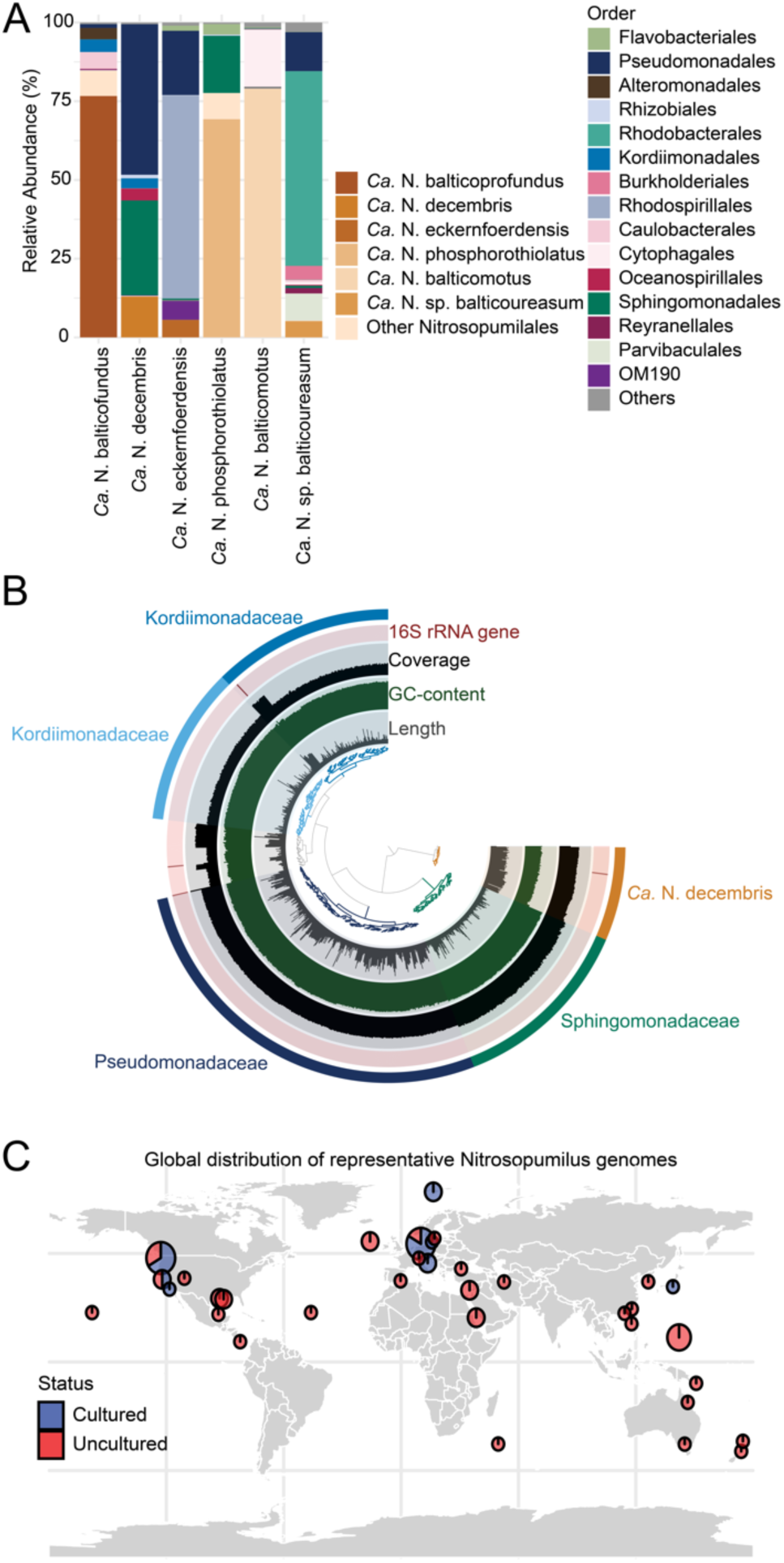
Enrichment cultures, genome binning, and distribution of AOA genomes. A) Microbial community composition analysis of different AOA strain enrichments. Nitrosopumilus was the only nitrifier detected and is shown at ASV level (brown shades), while other bacterial taxa are shown at the order level. *Ca.* N. phosphorothiolatus and *Ca.* N. balticomotus cultures show reduced bacterial diversity and proportions due to additional antibiotic treatment. B) Manual binning visualization of metagenomic assembly in Anvi’o, highlighting the distinct features of the archaeal bin including low GC-content and coverage patterns. C) Among 60 genomes analyzed in this study, 55 genomes are shown excluding four genomes without sampling location information.

While we were unable to enrich the dominant surface AOA strain from the southwestern Baltic Sea (ASV N. kielensis), we recovered a representative MAG through bulk metagenomic approaches. This MAG, which we name *Ca.* Nitrosopumilus sp. kielensis after its prevalence near Kiel, Germany, shares 98.5% ANI with a formally undescribed MAG from the Black Sea (GCA-14384445). This MAG’s ammonia monooxygenase alpha subunit (amoA) gene shows 98.6% nucleotide identity to HQ713535.1, which is the closest reference sequence to the dominant AOA (“aOTU1”) reported in surface water across both the northern (Öre Estuary) and southern Baltic Sea (Bay of Gdańsk)[39]. This high sequence similarity suggests that *Ca.* N. kielensis, or its close relatives, are widely distributed throughout Baltic Sea surface waters.

By mapping available Baltic Sea metagenomes against our comprehensive set of AOA reference genomes, distinct biogeographical patterns were revealed (Supplementary Fig. 5). In particular, analysis of shallow (<25 m) and deep (>25 m) samples showed that *Ca.* N. kielensis dominated the western Baltic Sea surface, while a closely related genome (92.1% ANI, 99.9% 16S rRNA gene identity; *Ca.* N. sp001437625 BACL13 MAG-121220-bin23) was predominantly detected in central and eastern Baltic Sea metagenomes. In the deep waters, the dominance of *Ca*.

Nitrosopumilus balticoprofundus was again confirmed by the mapping analysis. This analysis revealed further differences in the niches at the genome level. Similar mapping approaches for AOB and NOB yielded insufficient reads for robust observations, highlighting the need for comparable genomic efforts focusing on these groups.

### Genome-based trait conservation and variability of AOA

To understand the genomic diversity of Baltic Sea AOA in the context of global *Nitrosopumilus* diversity, we constructed a comprehensive phylogenomic tree incorporating genome sources, genomic characteristics, functional gene annotations and pangenomic analysis (Fig. 6). While Baltic Sea AOA were distributed across several clusters in the tree, most grouped with other brackish strains. All genomes contained core AOA genes essential for their lifestyle, including carbon fixation and ammonia oxidation pathways, or UV-damage repair (uvr) genes. In contrast, certain auxiliary functions previously observed in AOA -urease, motility, phosphorothioation, osmoregulation, and phosphate transport -were sparsely distributed among our Baltic Sea genomes. Among these, we found urease in *Ca.* N. balticoureasum, motility genes in *Ca.* N. balticomotus, phosphorothioation systems in *Ca.* N. phosphorothiolatus, and high-affinity phosphate-binding protein PstS in *Ca.* N. eckernfoerdensis and *Ca.* N. balticomotus. Notably, these traits showed no clear phylogenetic pattern in our analysis (Fig. 6). This is exemplified by two dominant surface strains that are closely related but differ in their metabolic capabilities: *Ca.* N. kielensis (western Baltic) lacks both pstS and urease, while N. sp001437625 BACL13 MAG-121220-bin23 (central Baltic) possesses both. While these metabolic differences might explain their distinct spatial distributions, testing this hypothesis requires additional environmental data, particularly urea concentrations.

**Fig. 6:**
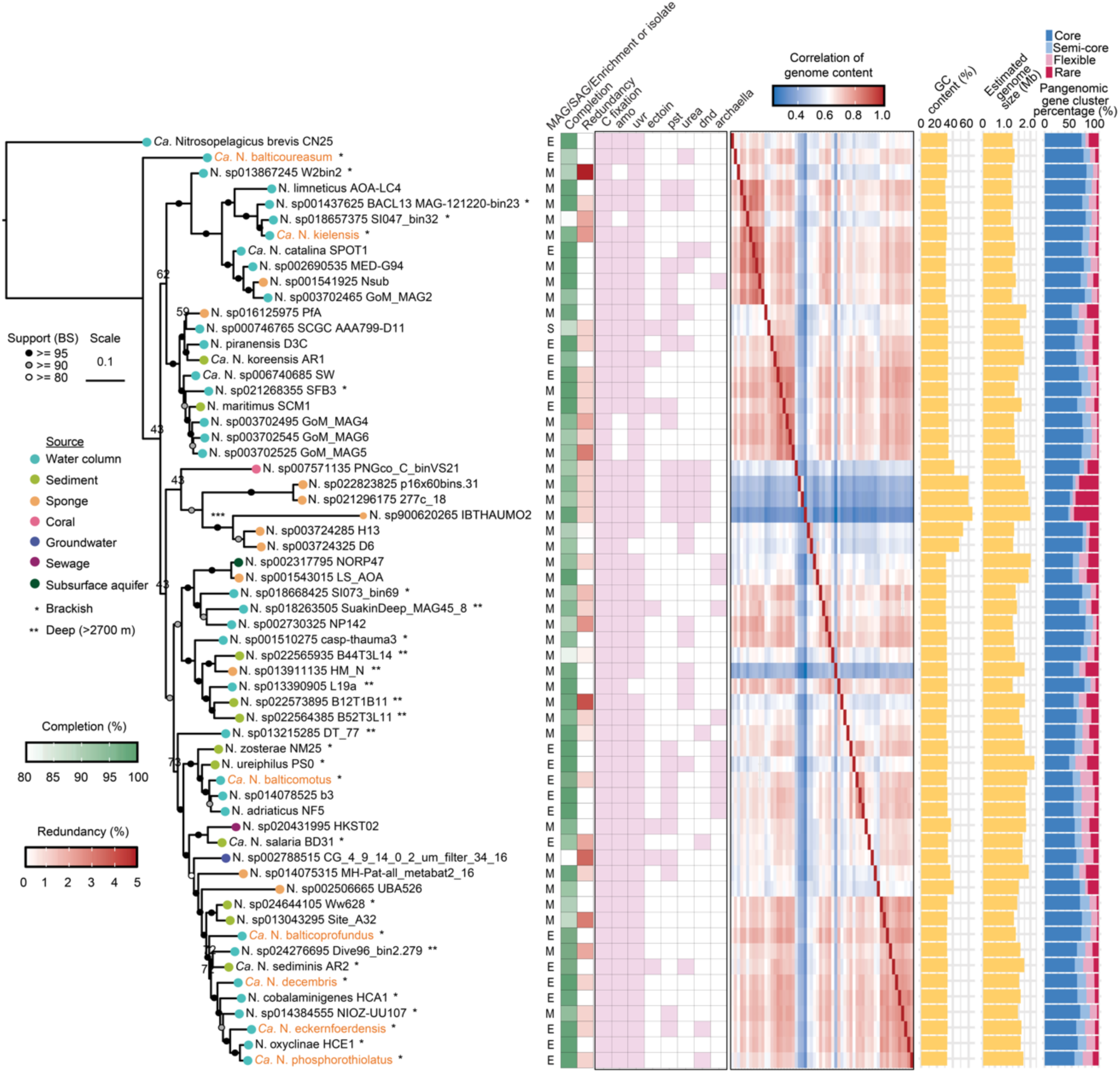
Comprehensive phylogenomic and functional analysis of 59 *Nitrosopumilus* genomes, with *Nitrosopelagicus brevis* as an outgroup. The phylogenomic tree was constructed based on single-copy core genes of Archaea, with genomes from this study highlighted in color. Genome sources are indicated by colored dots at branch tips, and habitat types (brackish/deep sea >2700m) are marked with asterisks next to the genome names. Basic genomic features (completion, redundancy, GC content, genome size, and genome type [MAG/SAG/enrichment or isolate]) are incorporated next to the tree. Selected functional gene sets are annotated in a presence/absence matrix (pink/white, respectively), where pink indicates the presence of >50% of genes in each category: carbon fixation (15 genes involved in 3HP/4HB cycle), ammonia oxidation (amo), UV repair (uvr), osmoregulation (ectoine), phosphate transport (pst), urease, DNA phosphorothioation (dnd), and motility (archaella). Pangenomic analysis revealed the distribution of gene clusters across genomes, categorized as core, semi-core, flexible, and rare genes, with their relative proportions shown for each genome. Genome content correlation was calculated based on gene presence/absence patterns in each gene cluster. *** indicates branches beyond this node were artificially shortened by half for visualization purposes. Detailed information about individual genes is provided in Supplementary Data 4.

Next, we used pangenomics to more fully examine gene conservation and flexibility across all *Nitrosopumilus*. Initially, a total of 14,118 gene clusters were identified. Among these, 7.84%, 2.34%, 14.15%, and 75.67% of clusters were considered “core”, “semi-core”, “flexible”, and “rare”, respectively (Supplementary Fig. 6). The core and semi-core genes represent an average of 75.3 ± 9.7% of the genes in each genome, while 13.3 ± 4.7% of the genes in a given genome were “flexible” and 11.4 ± 9.7% of the genes were “rare” (Fig. 6, rightmost panel). The substantial percentage of flexible and unique genes highlights the extensive genomic diversity within *Nitrosopumilus* both globally and within the Baltic Sea representatives. While most flexible and rare genes were un-annotated (30.3%, and 55% respectively), analysis of annotated genes by COG categories shows that genes involved in environmental interactions have higher proportions of variable genes (Fig. 7). For example, large proportions of genes in signal transduction mechanisms, Cell Wall/Membrane/Envelope Biogenesis, and Inorganic Ion Transport and Metabolism were categorized as semi-core, flexible, or rare within the pangenome.

**Fig. 7:**
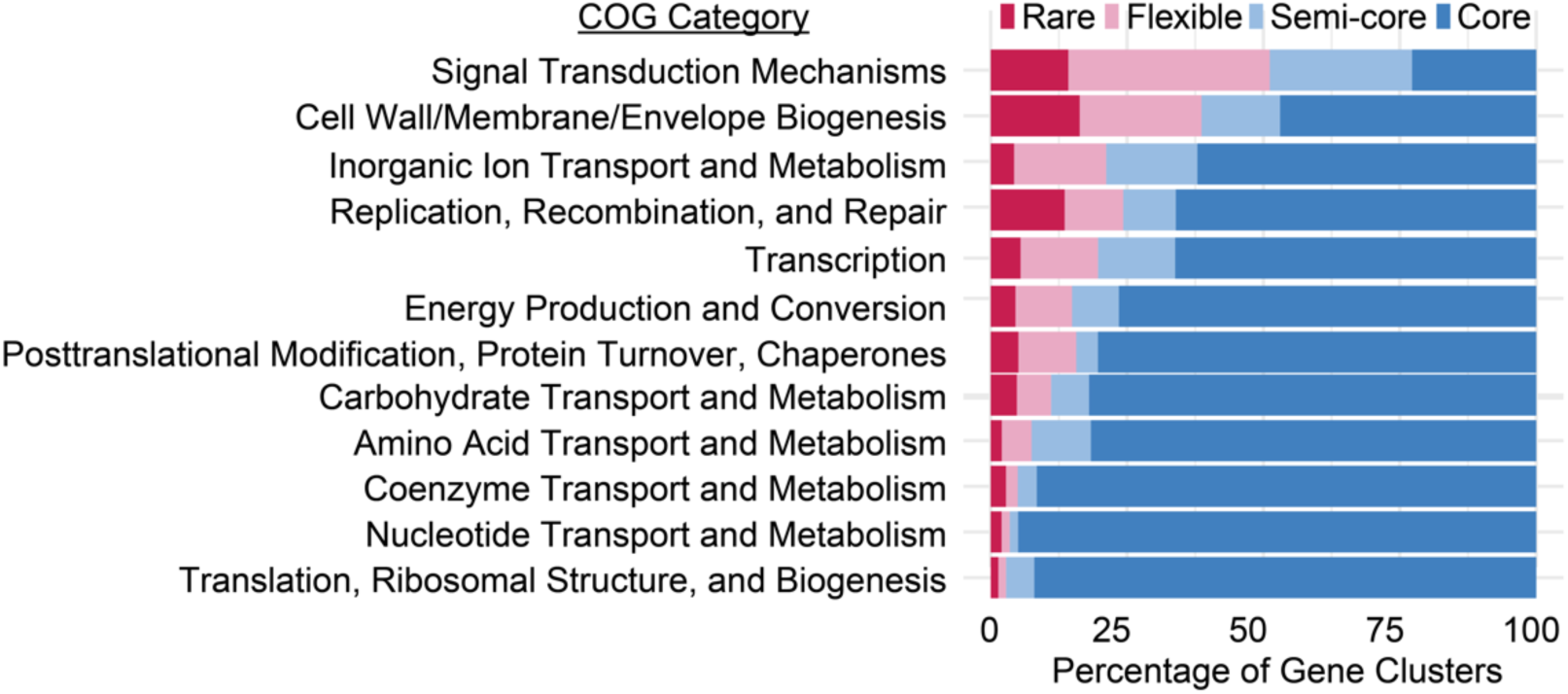
Distribution of COG20 functional categories across core, semi-core, flexible, and rare gene clusters in *Nitrosopumilus* representative genomes. Core and semi-core genes are predominantly associated with essential cellular functions, while flexible and rare genes show enrichment in environmental interaction processes, including signal transduction, cell wall/membrane biogenesis, and inorganic ion transport. Genes were excluded from the analysis if they were annotated as unknown function, had multiple category assignments, or belonged to categories with less than 2,000 genes.

Despite these many variable features, linking specific genomic traits to environmental adaptability appears complex for AOA, because of a variety of factors including 1.) many notable traits are not conserved across closely-related lineages, 2.) the overall pattern of gene presence and absence is largely driven by unannotated genes, and 3.) our understanding of the local and global distributions of AOA strains and candidate species remains limited. Systematically assessing distributions beyond the Baltic Sea for all AOA, may help further address this important question.

Further work utilizing methods such as transcriptomics and proteomics would offer a further understanding of how gene potential functions are expressed and contribute to each organism’s distributions, for example through gene networks and activities in response to environmental variation. Additionally, the key to these strain differentiations may involve enzyme kinetics or other attributes that cannot be readily predicted from genomic data alone. Our research and novel genomes provide a foundation for future work to help uncover how these organisms adapt and thrive in the changing conditions of the Baltic Sea.

## CONCLUSIONS

By integrating time-series and distributional data with genomic analysis, this study deepens our understanding of the realized niche and potential drivers of nitrification in a rapidly changing ecosystem, contributing to our knowledge of nitrifier ecology. Our results revealed fine-grained niche-partitioning across the Baltic Sea, with late-fall to winter-time increase in nitrifiers in surface waters in the southwest Baltic Sea. The seasonal dynamics in the surface were tightly coupled to geochemistry in the water column, where nitrite accumulated early in the winter, followed by nitrate accumulating slower until the spring bloom. Further temporal and spatial sampling, along with coupled environmental measurements and rate measurements (such as nitrification and carbon fixation), will help to advance an understanding of the consistency and/or variability of the inter- and intra-annual variability of the nitrifiers, and the potential drivers and implications. Meanwhile, the enrichment of novel AOA strains from the Baltic Sea revealed that strains share most of their genes, yet each genome contained large amounts of flexibility or novelty, particularly in genes involved in environmental interactions. However, connecting specific gene repertoires to strain-specific to environmental distributions remained elusive. Together, our time-series and distributional study combined with novel genomes sets the stage for more detailed and mechanistic exploration into the bottom-up and top-down factors controlling the ecology and evolution of nitrifiers in the Baltic Sea and ultimately towards other environments such as the deep ocean.

## Author Contributions

SK analyzed amplicon, genomic, and environmental data. DMN analyzed Baltic Sea-wide metagenomes. SK and DMN enriched and maintained AOA enrichments. SK extracted and sequenced AOA enrichments. ED established protocols for DNA extraction and sequencing in the lab and performed the molecular work for field samples with support from DMN. SK and DMN wrote the paper with edits from ED. DMN secured funding for the research, via Helmholtz Young Investigator Grant and Deutsche Forschungsgemeinschaft grant NE 2754/1-1.

## Data Availability

Previously unpublished environmental data utilized in the manuscript are available via Supplementary Data 1. Metagenome and amplicon short read datasets are available via NCBI SRA (upon publication). Newly described AOA genomes are available via NCBI Accessions (upon publication). Alignments and phylogenetic trees are available via Figshare (DOI:). Prior to publication, all data available upon request to the authors.

## Acknowledgments

We thank all those involved in our time-series projects, including Alyzza Calyag, Karolin Eisenschmid, Selina Ernst, Clara Fenge, Marjan Ghotbi, Maarten Kanitz, Ana Belén Kuhlmann, Lara Schroeder, and all other members of the Marine Microbial Ecology group at GEOMAR. We thank SE for helping enrich AOA enrichments. We thank MG who shared an observation of high AOA abundance in the Kiel Fjord prior to our study. We thank Jan Muschiol and Kerstin Petersen for general support. We thank the chief scientists who facilitated the samplings from the Baltic Sea, Jan Dierking (AL571) and Felix Mittermayer (AL580), as well as to all participants in the cruises. We thank Hermann Bange and Kastriot Qelaj for organizing monthly Boknis Eck cruises. Also, the valuable environmental data was from KIMOCC thanks to Claas Hiebenthal and Frank Melzer.

## Protologues for new *Candidatus* species identified from enrichments and metagenomics for five novel *Nitrososopumilus* species

### Description of *Candidatus* Nitrososopumilus balticoprofundus sp. nov

*Candidatus* Nitrososopumilus balticoprofundus (bal.ti.co.pro.fun’dus. N.L. masc./fem. adj. balticus of or pertaining to the Baltic; L. adj. profundus deep; N.L. balticoprofundus predominance in deep Baltic Sea waters).

An archaeal species identified by enrichment-enabled genomics. This genome originated from a sample from the Baltic Sea, Bornholm Basin, from a depth of 63 m, and was enriched in autoclaved Baltic Sea water amended with ammonium, phosphate, trace metals, and antibiotics. The enrichment was collected on a 0.2 µm filter, DNA extracted, sequenced by shotgun metagenomics with Illumina Sequencing, assembled with spades, and manually curated with Anvi’o. This species shows less than 95% Average nucleotide identity to the type *Nitrosopumilus* genome, and all representative genomes in the GTDB database. The phylogenetic position was analyzed using reference genomes from the GTDB database. The genome is estimated to be 97.09% complete and 0% contaminated. The GC Content of the genome is 32.4%, genome size is 1.34 Mb, with 40 contigs. The genome is the numerically dominant archaeal species found in the deep Baltic Sea, according to mapping of metagenomes at 95% ANI.

### Description of *Candidatus* Nitrososopumilus eckernfoerdensis sp. nov

*Candidatus* eckernfoerdensis (ec.kern.foer.den’sis. N.L. masc./fem. adj. eckernfoerdensis referring to Eckernförde, a city in Germany near the location in the Baltic Sea from which the water for enrichment was collected).

An archaeal species identified by enrichment-enabled genomics. This genome originated from a sample from the southwest Baltic Sea, Kiel Bight (Boknis Eck sampling location), from a depth of 5 m, and was enriched in autoclaved Baltic Sea water amended with ammonium, phosphate, trace metals, and antibiotics. The enrichment was collected on a 0.2 µm filter, DNA extracted, sequenced by shotgun metagenomics with Illumina Sequencing, assembled with spades, and manually curated with Anvi’o. This species shows less than 95% Average nucleotide identity to the type *Nitrosopumilus* genome, and all representative genomes in the GTDB database. The phylogenetic position was analyzed using reference genomes from the GTDB database. The genome is estimated to be 99.03% complete and 0.97% contaminated. The GC Content of the genome is 33.4%, genome size is 1.62 Mb, with 12 contigs. The genome is the numerically rare archaeal species found in the southwestern Baltic Sea (Kiel Bight), according to mapping of metagenomes at 95% ANI and 16S rRNA sequence analysis

### Description of *Candidatus* Nitrososopumilus balticomotus sp. nov

*Candidatus* Nitrososopumilus balticomotus (bal.ti.co.mo’tus. N.L. masc./fem. adj. balticus of or pertaining to the Baltic; L. adj. motus moved; N.L. balticomotus referring to referring to the presence of an archaellum gene in its genome, a discerning feature of the genome.).

An archaeal species identified by enrichment-enabled genomics. This genome originated from a sample from the southwest Baltic Sea, Kiel Bight (Kiel Fjord sampling location), from a depth of 1 m, and was enriched in filter-sterilized (0.2 µm) Baltic Sea water amended with ammonium, phosphate, trace metals, and antibiotics. The enrichment was collected on a 0.2 µm filter, DNA extracted, sequenced by shotgun metagenomics with Illumina Sequencing, assembled with spades, and manually curated with Anvi’o. This species shows less than 95% Average nucleotide identity to the type *Nitrosopumilus* genome, and all representative genomes in the GTDB database. The phylogenetic position was analyzed using reference genomes from the GTDB database. The genome is estimated to be 97.09% complete and 0.97% contaminated. The GC Content of the genome is 32.6%, genome size is 1.83 Mb, with 36 contigs. The genome is the numerically rare archaeal species found in the southwestern Baltic Sea (Kiel Bight), according to mapping of metagenomes at 95% ANI and 16S rRNA sequence analysis

### Description of *Candidatus* Nitrososopumilus decembris sp. nov

*Candidatus* Nitrososopumilus decembris (de.cem’bris. L. adj. decembris pertaining to December; N.L. decembris indicating the month from which the water for enrichment was collected from Baltic Sea surface waters).

An archaeal species identified by enrichment-enabled genomics. This genome originated from a sample from the southwest Baltic Sea, Kiel Bight (Kiel Fjord sampling location), from a depth of 1 m, and was enriched in filter-sterilized (0.2 µm) Baltic Sea water amended with ammonium, phosphate, trace metals, and antibiotics. The enrichment was collected on a 0.2 µm filter, DNA extracted, sequenced by shotgun metagenomics with Illumina Sequencing, assembled with spades, and manually curated with Anvi’o. This species shows less than 95% Average nucleotide identity to the type *Nitrosopumilus* genome, and all representative genomes in the GTDB database. The phylogenetic position was analyzed using reference genomes from the GTDB database. The genome is estimated to be 100% complete and 0.97% contaminated. The GC Content of the genome is 33.3%, genome size is 1.5 Mb, with 11 contigs. The genome is the numerically rare archaeal species found in the southwestern Baltic Sea (Kiel Bight), according to mapping of metagenomes at 95% ANI and 16S rRNA sequence analysis

### Description of *Candidatus* Nitrososopumilus phosphorothiolatus sp. nov

*Candidatus* Nitrososopumilus phosphorothiolatus (phos.pho.ro.thi.o.la’tus. N.L. neut. n. phosphorus relating to phosphate; G. n. thiol relating to sulfur compounds; N.L. phosphorothiolatus referring to the presence of a phosphothioation pathway in its genome, a discerning feature of the genome).

An archaeal species identified by enrichment-enabled genomics. This genome originated from a sample from the southwest Baltic Sea, Kiel Bight (Kiel Fjord sampling location), from a depth of 1m, and was enriched in filter-sterilized (0.2 µm) Baltic Sea water amended with ammonium, phosphate, trace metals, and antibiotics. The enrichment was collected on a 0.2 µm filter, DNA extracted, sequenced by shotgun metagenomics with Illumina Sequencing, assembled with spades, and manually curated with Anvi’o. This species shows less than 95% Average nucleotide identity to the type *Nitrosopumilus* genome, and all representative genomes in the GTDB database. The phylogenetic position was analyzed using reference genomes from the GTDB database. The genome is estimated to be 99.51% complete and 0.97% contaminated. The GC Content of the genome is 32.8%, genome size is 1.7 Mb, with 11 contigs. The genome is the numerically rare archaeal species found in the southwestern Baltic Sea (Kiel Bight), according to mapping of metagenomes at 95% ANI and 16S rRNA sequence analysis.

### Description of *Candidatus* Nitrosopumilus balticoureasum sp. nov

*Candidatus* Nitrosopumilus balticoureasum (bal.ti.co.u.re’a.sum. N.L. masc./fem. adj. balticus of or pertaining to the Baltic; L. adj. ureasum referring to urea; N.L. balticoureasum referring to the presence of a urease gene in its genome, a discerning feature of the genome)

An archaeal species identified by enrichment-enabled genomics. This genome originated from a sample from the southwest Baltic Sea, Kiel Bight (Kiel Fjord sampling location), from a depth of 1m, and was enriched in filter-sterilized (0.2 µm) Baltic Sea water amended with ammonium, phosphate, trace metals, and antibiotics. The enrichment was collected on a 0.2 µm filter, DNA extracted, sequenced by shotgun metagenomics with Illumina Sequencing, assembled with spades, and manually curated with Anvi’o. This species is 95.33% average nucleotide identity to the closest representative genome in the GTDB database (sp008080815). This closest relative is a strain identified by metagenomics coastal Sapelo Island, Georgia, USA, which to-date is not formally described, and has only the species place-holding number (sp008080815). The phylogenetic position was analyzed using reference genomes from the GTDB database. The genome is estimated to be 89.32% complete and 0% contaminated. The GC Content of the genome is 32.9%, genome size is 1.19 Mb, with 75 contigs.

### Description of *Candidatus* Nitrosopumilus kielensis sp. nov

*Candidatus* Nitrosopumilus kielensis (kiel.en’sis. N.L. masc./fem. adj. kielensis, derived from Kiel, a city in Germany associated with the location of study).

An archaeal species identified by metagenomics. This genome originated from a sample from the southwest Baltic Sea, Kiel Bight, Boknis Eck sampling location from a depth of 25 meters in February 2022. The seawater was collected on a 0.2 µm filter, DNA extracted, sequenced by shotgun metagenomics with Illumina Sequencing, assembled with spades, and manually curated with Anvi’o. This species was 98.5% average nucleotide identity to the closest representative genome in the GTDB database (sp014384555). This closest relative is also a strain identified by metagenomics from the Black Sea, which to-date was not formally described, and has only a species place-holding number (sp014384555). The phylogenetic position was analyzed using reference genomes from the GTDB database. The genome is estimated to be 96.12% complete and 2.27% contaminated. The GC Content of the genome is 32.5%, genome size is 1.26 Mb, with 50 contigs. The genome is the numerically dominant archaeal species found in the southwestern Baltic Sea (Kiel Bight), according to mapping of metagenomes at 95% ANI.

