## Supplementary Figures and Legends for "Basin-scale dynamics and enrichment-enabled genomics of marine nitrifiers: seasonality, niches, interactions, and genomic uniqueness"

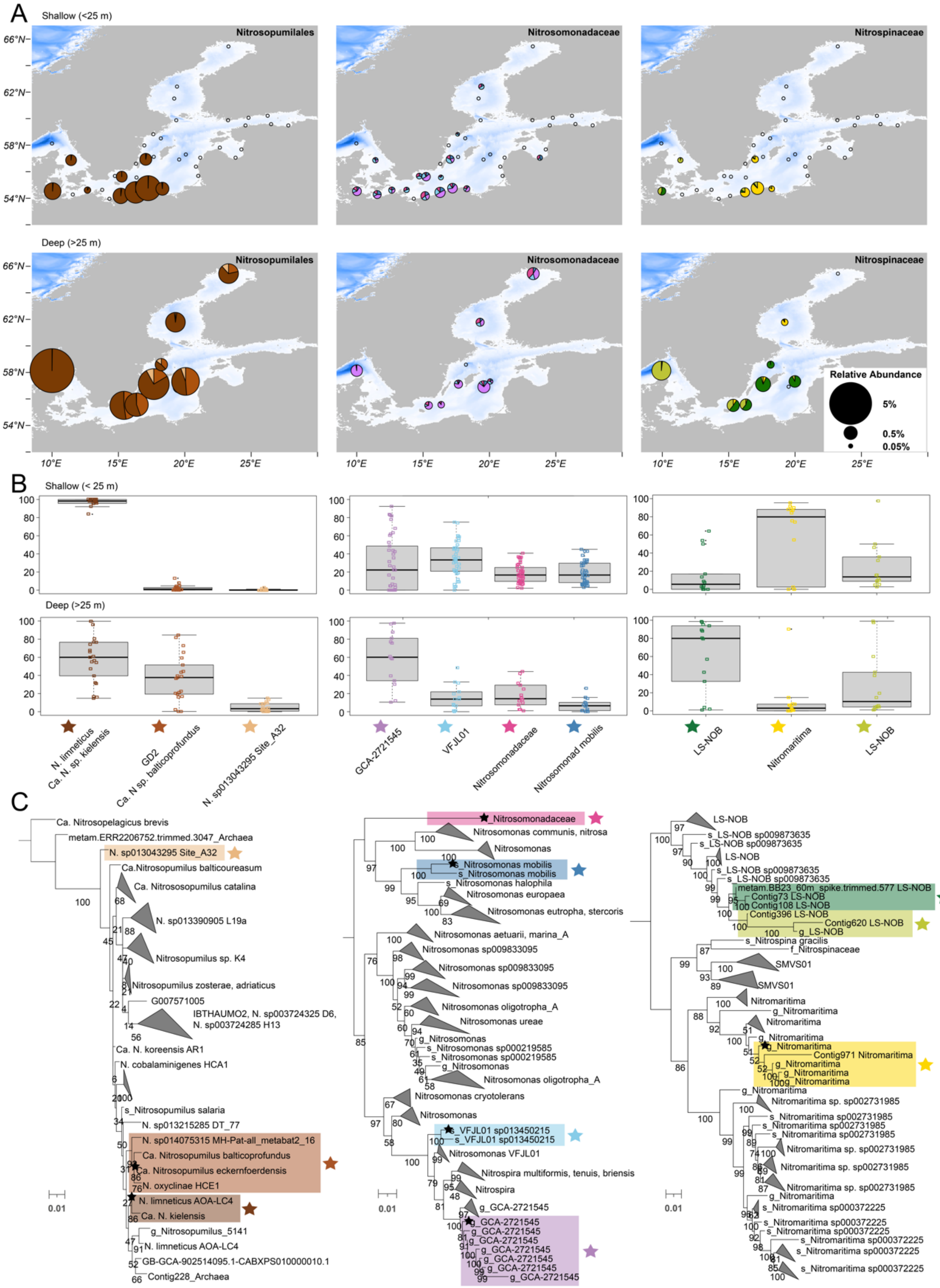


**S Fig 1:** Distributions of nitrifiers and phylotypes across the entire Baltic Sea. The figure is based on the same data as Figure 1, but includes a breakdown of the different phylotypes observed based on phylogenetic placement. A) The pie size indicates the average relative abundance of the given family at each location (all samples within 1° of latitude and longitude are averaged) with the respective phylotypes indicated by color. Upper panels show surface samples (0-25 m), and the lower panels show deep samples (>25 m). Open black circles indicate no detection. The background colors of the Baltic Sea (whites and blues) indicate water column depth. B) Summary box-and-whisker plot illustrating the proportion of each phylotype across all samples for a given depth range. C) The phylogenetic trees are as described in the methods. Collapsed branches are phylotypes that were not significantly detected or detected at all in the data.

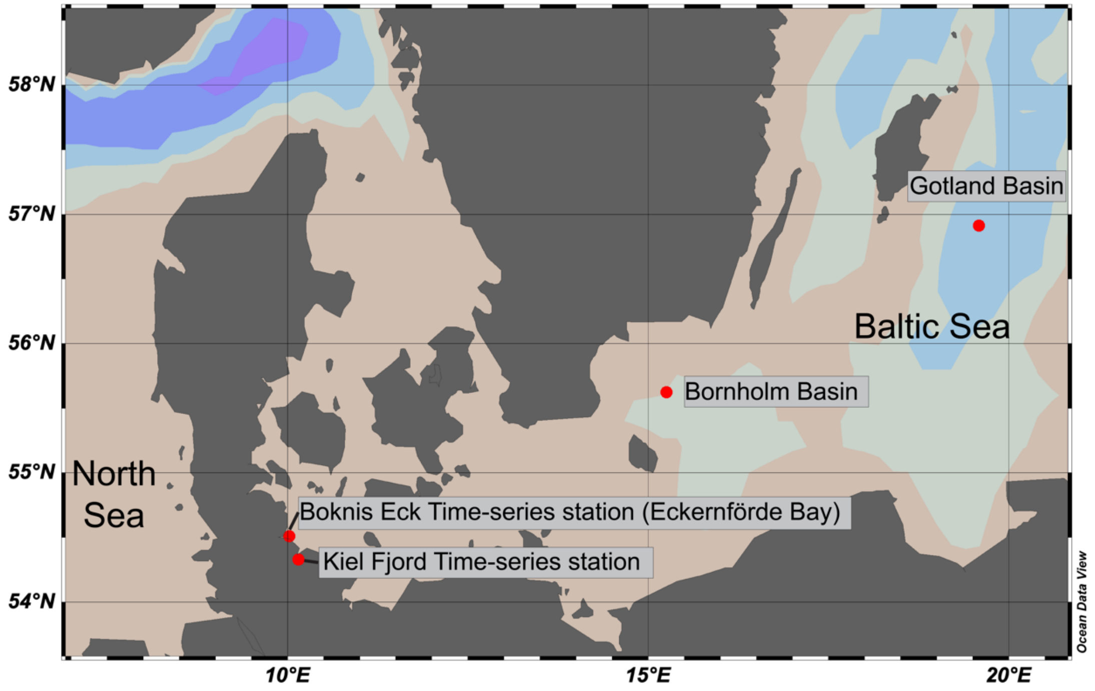


**S. Fig 2:** Map of sampling locations from this study. Sampling sites include the Kiel Fjord Time-series station (twice-weekly sampling from October 2021 to May 2023), Boknis Eck Time-series station in Eckernförde Bay (monthly sampling from January 2022 to June 2023), and broader Baltic Sea region sampling sites at Bornholm Basin and Gotland Basin (September 2022). Previously published metagenomic data sampling locations are not included in this map.

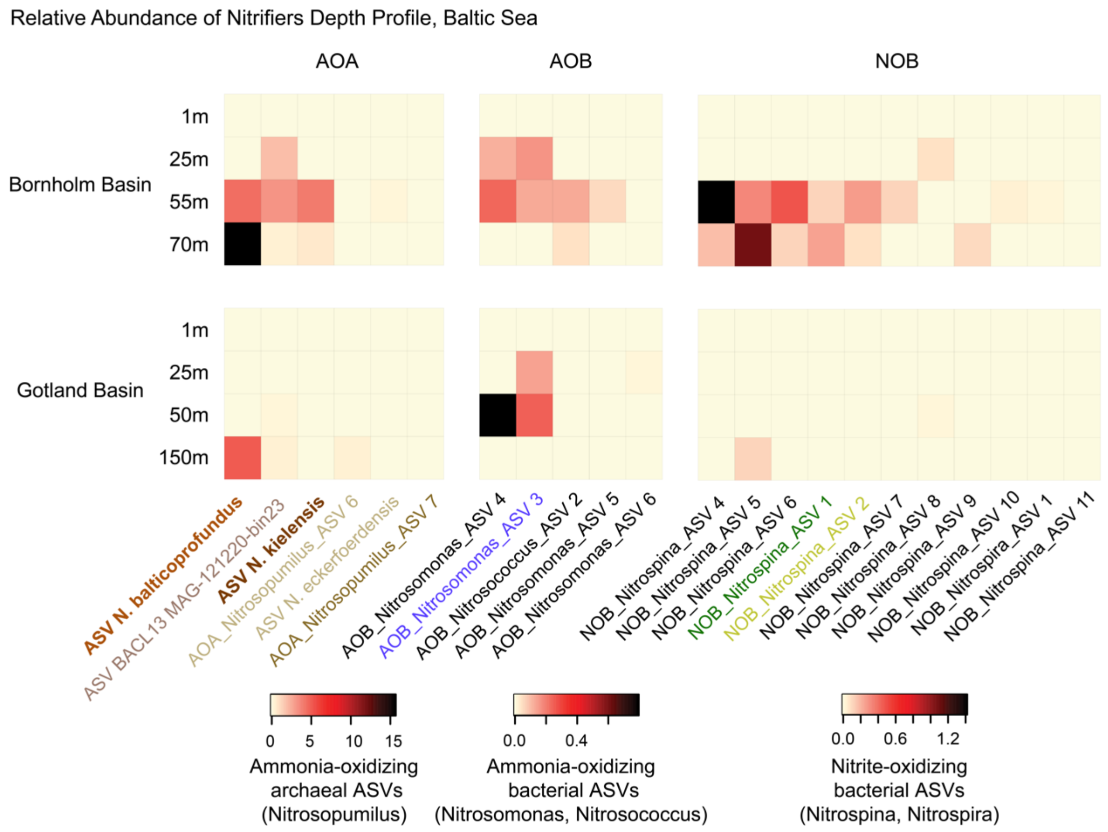


**S. Fig 3:** Nitrifier depth distributions at two sampling locations in the Baltic Sea (Bornholm and Gotland Basins). Sampling in early September revealed nitrifier communities were barely detected in surface waters but maintained substantial populations in deeper waters (>25 m). AOA dominated the nitrifier communities, particularly ASV N. balticoprofundus, which showed highest relative abundance in deep waters at both sites. While AOA and NOB maintained substantial populations in deeper waters, AOB abundance was limited to intermediate depths. The depth distribution patterns, particularly the prevalence of ASV N. balticoprofundus and NOB_Nitrosopina_ASV5 and absence of AOB in the bottom water, suggest differential adaptation of nitrifier to oxygen-limited conditions

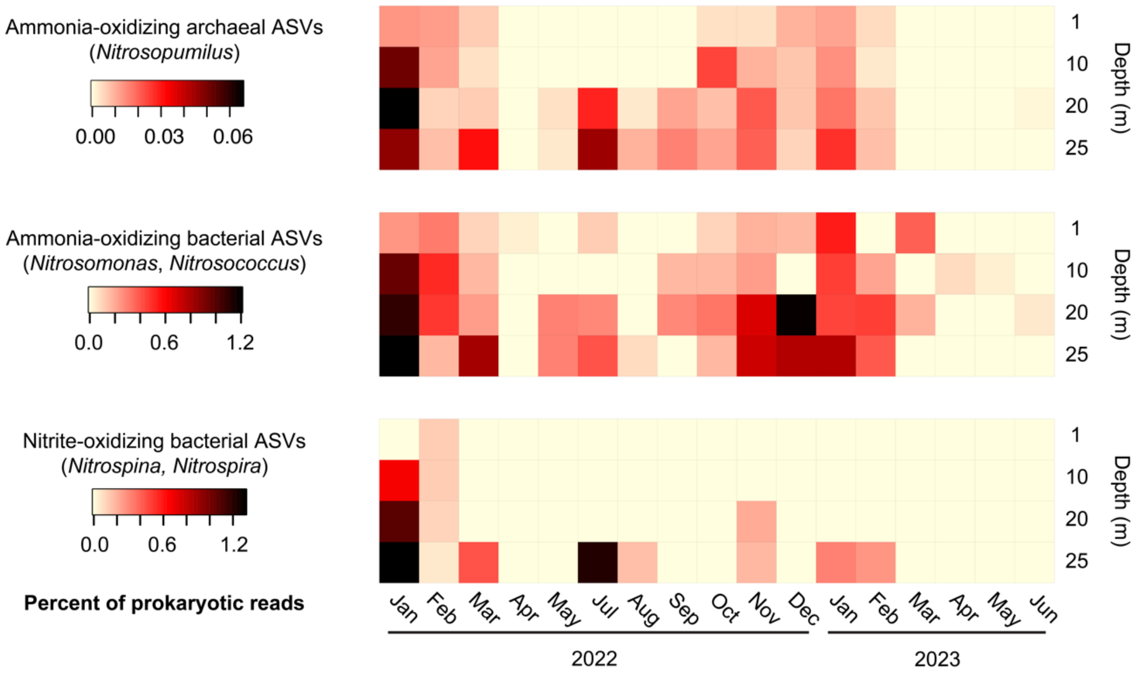


**S. Fig 4:** Nitrifier depth and temporal dynamics at Boknis Eck Time-series station located in the southwest Baltic Sea. In order to contextualize the surface dynamics in the Kiel Fjord surface waters, we also examined nitrifier dynamics spanning the same time-period from surface to seafloor (1m to 25m) in the southwest Baltic Sea at the Boknis Eck time-series station. This depth-integrated monthly time-series demonstrated nitrifiers were also abundant across the whole water column during the winter, but were even more abundant at deeper depths (20 m and 25m), compared with surface waters. During summer months, nitrifier populations persisted in the deeper waters (mainly 20-25 m), showing occasional increases in abundance.

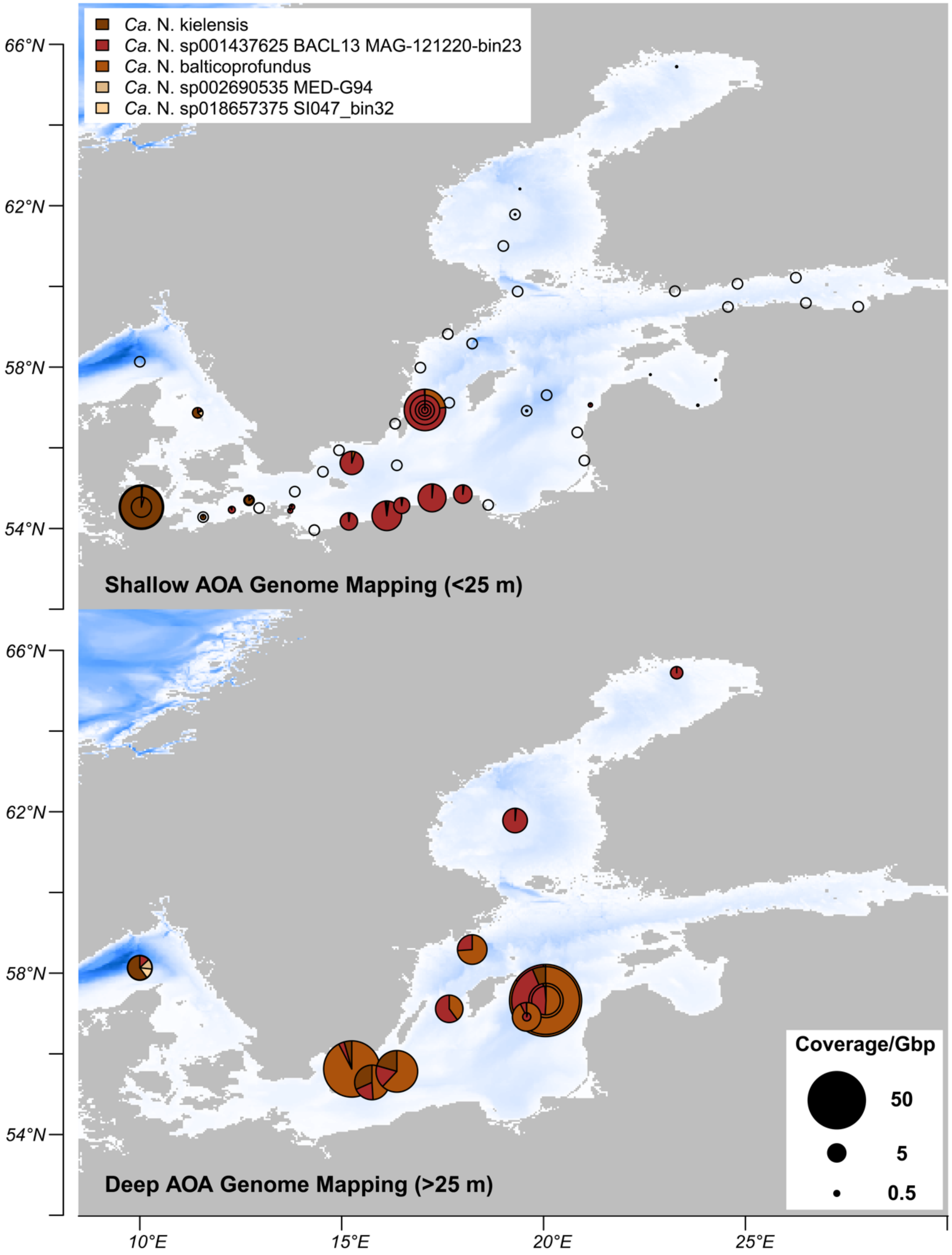


**S. Fig 5:** Coverage of AOA genomes from metagenomes across the entire Baltic Sea. Only genomes that cumulatively had greater than 25x coverage across all samples are displayed.

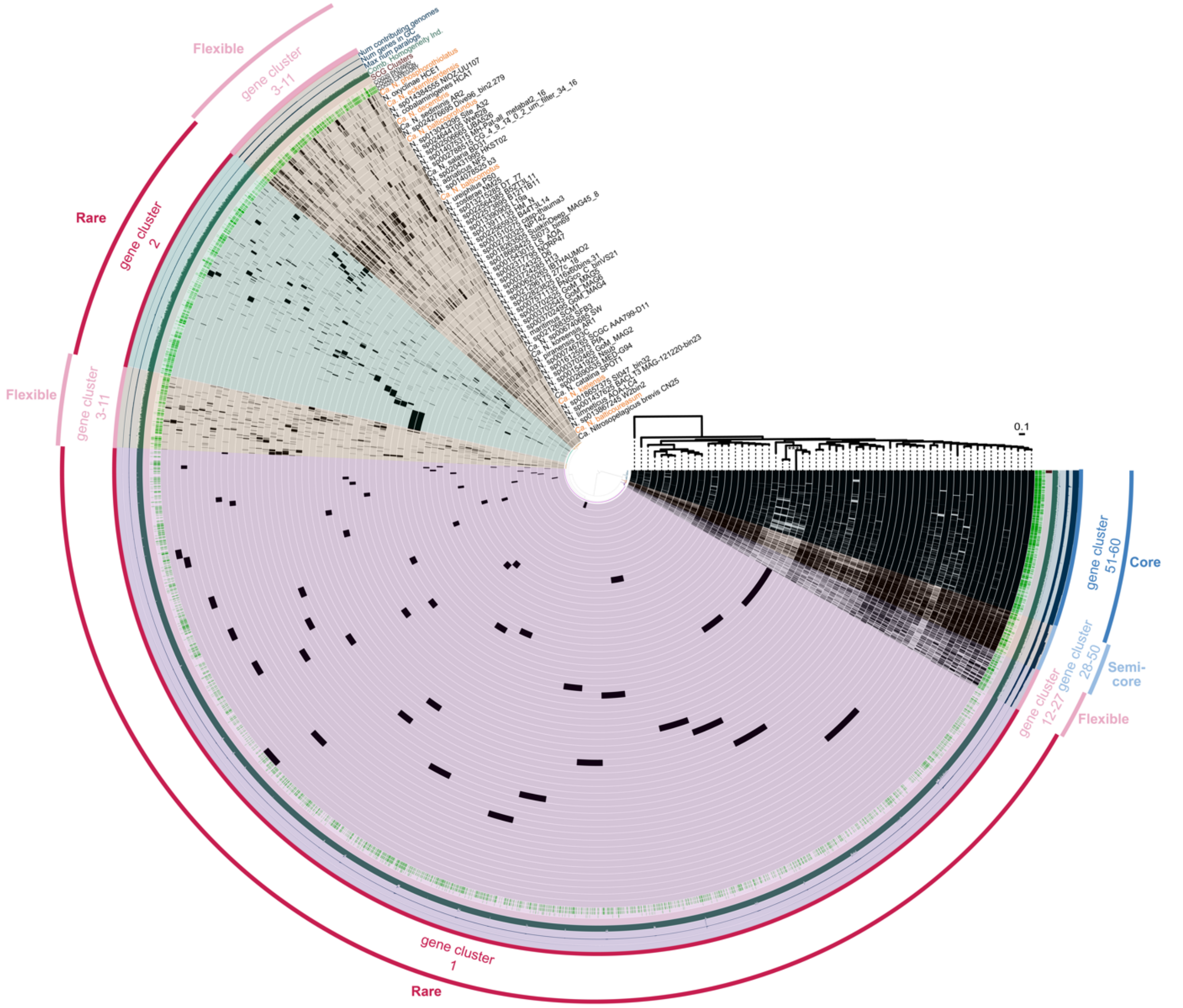


**S. Fig 6:** The Pan-genome of all AOA representative genomes used in this study. Each bar in the circularized plot represents a gene cluster and is filled if the corresponding genome contributes a gene to that cluster. The outermost layer classifies gene clusters into categories (core, semi-core, etc.), where numerical ranges (51-60, 28-50, etc.) indicate how many genomes contribute to a gene cluster for it to be assigned to a given category. The inner dendrogram clusters homologous gene families based on their presence/absence patterns across genomes. Genome order follows the phylogenomic tree (shown middle-right), where branches marked with *** represent deep-branched sponge-associated genomes. (as in Fig. 6). The outer layers indicate 1.) the number of genomes contributing to each gene cluster, 2.) number genes in Gene Clusters, 3.)

maximum number of paralogs, 4.) combined homogeneity index; indicative for similarity of sequences within a gene cluster, 5.) SCG Clusters; single copy gene clusters, 6.) COG20 PATHWAY, 7.) COG20 FUNCTION, 8.) COG20 CATEGORY. That is, 6.), 7.), and 8.) indicate where a gene has an annotation in the COG database. Most of the rare genes lack COG annotations, while SCGs typically have annotations.
